# Ancestral contributions to contemporary European complex traits

**DOI:** 10.1101/2021.08.03.454888

**Authors:** Davide Marnetto, Vasili Pankratov, Mayukh Mondal, Francesco Montinaro, Katri Pärna, Leonardo Vallini, Ludovica Molinaro, Lehti Saag, Liisa Loog, Sara Montagnese, Rodolfo Costa, Mait Metspalu, Anders Eriksson, Luca Pagani

**Author notes:** Corresponding Authors (DM); (LP).

## Abstract

The contemporary European genetic makeup formed in the last 8000 years as the combination of three main genetic components: the local Western Hunter-Gatherers, the incoming Neolithic Farmers from Anatolia and the Bronze Age component from the Pontic Steppes. When meeting into the post-Neolithic European environment, the genetic variants accumulated during their three distinct evolutionary histories mixed and came into contact with new environmental challenges.

Here we investigate how this genetic legacy reflects on the complex trait landscape of contemporary European populations, using the Estonian Biobank as a case study.

For the first time we directly connect the phenotypic information available from biobank samples with the genetic similarity to these ancestral groups, both at a genome-wide level and focusing on genomic regions associated with each of the 27 complex traits we investigated. We also found SNPs connected to pigmentation, cholesterol, sleep, diastolic blood pressure, and body mass index (BMI) to show signals of selection following the post Neolithic admixture events. We recapitulate existing knowledge about pigmentation traits, corroborate the connection between Steppe ancestry and height and highlight novel associations. Among others, we report the contribution of Hunter Gatherer ancestry towards high BMI and low blood cholesterol levels.

Our results show that the ancient components that form the contemporary European genome were differentiated enough to contribute ancestry-specific signatures to the phenotypic variability displayed by contemporary individuals in at least 11 out of 27 of the complex traits investigated here.

## 1 Introduction

Since its origins, ancient human genetics showed that the current European genetic landscape formed only recently, in the last 8000 years, as the combination of three main genetic components: 1) the local Western Hunter-Gatherers (WHG), 2) the incoming Neolithic Farmers from the Near East (Anatolia_N) and 3) the Bronze Age component from the Pontic Steppes, often identified with the Yamnaya culture (Yamnaya)^1–3^. As a result, any modern European population is a combination of at least these three components, in variable proportions depending on its particular genetic history and geographic location. Before their arrival in Europe, the ancestors of these three components evolved in different areas and environments for thousands of years, hence differentiating through neutral genetic drift but also adapting to the different climatic, nutritional and pathogenic conditions. When coming together into the post-Neolithic European environment, the genetic variants accumulated during their three distinct evolutionary histories admixed and came into contact with new environmental challenges.

Previous research efforts have indeed characterized evolutionary events specific to these populations which putatively affected their phenotype and appearance, through the tracking of few highly characterized SNPs^4–6^ or polygenic scores^7,8^. Nevertheless, while the first approach is limited in the number of variants analyzed and largely blind with regards to complex polygenic traits, the second builds on population-dependent effect sizes^9,10^ estimated in Genome Wide Association Studies (GWAS). Such summary statistics, especially in their genome-wide aggregations, may lead to directional bias and lower predictive accuracy in populations different to the one where the GWAS study was performed^11–15^ and have sometimes led to ambiguous results about polygenic adaptation^16–19^.

Here we capitalize on the Estonian Biobank by measuring the relative genetic distance of contemporary individuals to a given ancestry and associating it with their phenotype, thus measuring the influence of these ancient genetic sources on the complex traits distribution of contemporary Europeans. By connecting directly the phenotypic information with the genetic similarity to these ancestral groups we avoid the step of producing and interpreting association summary statistics which might compress information or produce the spurious results mentioned above. For the same reason, our conclusions are applicable to contemporary individuals of European ancestry, where the phenotypes were collected. Conversely, using them to extrapolate features of ancient populations, although tempting, should be done with caution due to the interaction of their genetic legacy with a radically different lifestyle and environment.

We started by selecting 27 complex traits of interest, for which we have sufficient data in the Estonian Biobank, a collection of samples from a relatively homogeneous European population which is among the ones with the highest fraction of remnant Hunter Gatherer genomic component and additionally includes a Siberian (Siberia) component associated with Iron Age movements^20,21^. In order to associate a phenotype to the contribution of a specific ancient European ancestry we introduce *covA*, the covariance between allele frequencies in contemporary individuals and a given ancestral population with respect to the contemporary and ancient average frequencies (see Methods and Supplementary Notes for further details). We computed *covA* for each pair of Estonian individuals and ancestries, defined models in which each trait is predicted by *covA* of a specific ancestry and used them to elucidate ancestry/trait associations. We refined our approach by focusing on the ancestry similarity patterns in genomic regions potentially connected to each trait according to GWAS catalog^22^. Based only on the SNPs contained in such regions, we then measured *covA* as above and used it as a predictor to model traits, also in comparison with random genomic sets with matching size. Finally we set out to independently analyze if those regions that are associated with the genetic contribution of a specific ancestry also experienced a post-admixture selective pressure.

## 2 Results

### 2.1 *covA* measures similarity with ancestral groups

We computed *covA* for each pair of Estonian individuals and ancestries among WHG, Anato-li_N, Yamnaya and Siberia using manually curated and other ancient individuals shortlisted by genetic and chronological proximity (see Methods and Table S1). By observing *covA* joint distributions in Figure 1a,b (see Figure S1 for all combinations) we can see that as expected, *covA*s calculated on the various ancestries are strongly interdependent, mainly because they include as term the average ancestral frequency and partly because of varying grades of similarity among the ancestries for historical demographic reasons. In particular they tend to be negatively correlated except for *covA* for Yamnaya being associated with *covA* for WHG, reflecting complex demographic relationships between the two, due to WHG-like Eastern Hunter Gatherer ancestry presence in Yamnaya^2,3,23^. Even if by European standards Estonia can be considered relatively genetically uniform, as recently shown in Pankratov *et al*. [24] the south-eastern inland counties tend to have higher haplotype sharing with Latvians, Lithuanians and Russians compared with the rest of the country, and especially the northern coast: this is reflected by median *covA* for WHG being higher in those Estonian counties, see Figure 1c. Conversely, as shown by median *covA* for Siberia in Figure 1c, the Siberian ancestry seems to be more abundant in the north-east, consistently with Finnish ancestry shown in Pankratov *et al*. [24]. Yamnaya and Anatolia_N *covA*s are instead more evenly distributed (Figure S2).

**Figure 1:**
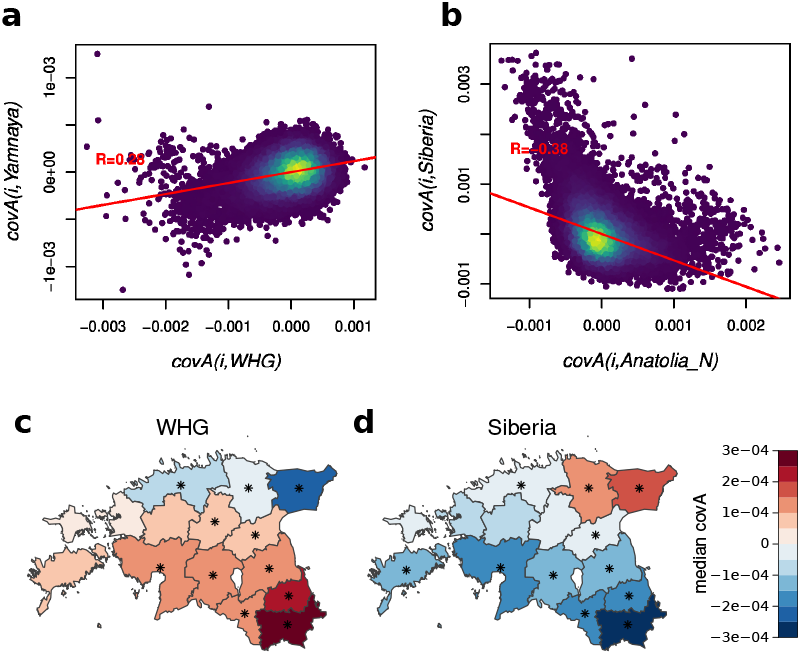
*covA* distributions. **a,b** *covA* joint distributions for two ancestry couplings. Each dot is an individual, dots in denser areas are lighter. The red line shows a linear regression, with its R coefficient. **c,d** *covA*s for WHG and Siberia across Estonian counties. Color indicates median *covA* computed in each county, with the sign reflecting excess or lack of a given ancestry, while asterisks indicate those counties for which the *covA* distribution is significantly different than the rest of Estonia (two-tailed Wilcoxon-Mann-Whitney test, *p* ≤ 0.001)

### 2.2 Connecting complex trait variation to genome-wide ancestry similarity

We examined 27 complex traits (31 if considering pigmentation variants) which were corrected and adjusted for covariates (including sex, age, genotyping platform and others, see Table S2), and expecting varying degrees of influence from genome-wide ancestry depending on their heritability as captured by our dataset(Figure 2a).

**Figure 2:**
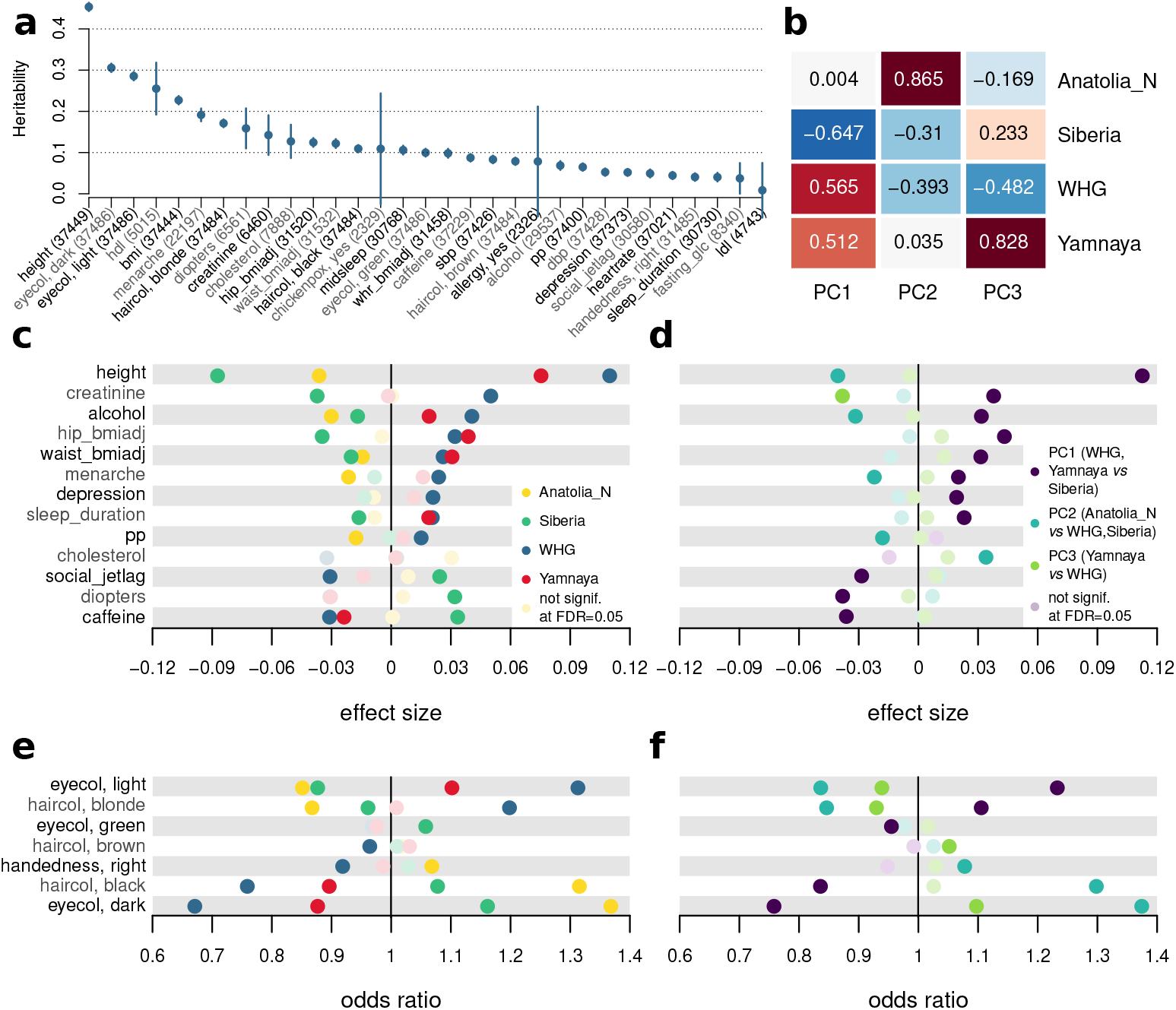
Genome-wide ancestry-trait associations. **a** All traits analyzed and their estimated heritability after covariate adjustment. Numbers in parentheses indicate the number of unrelated samples for which phenotypic information was available for each trait. **b** Loading matrix for genome wide *covA*s and their PCs. PCs can be interpreted as axes defined by 2 or 3 *covA*s. **c-f** Genome-wide *covA* or *covA*-based PCs estimated coefficients for traits which have at least one significant PC coefficient. *β* for continuous **c,d** and Odds Ratios (OR) for categorical **e,f** traits. Pastel dots are deemed not significant at Benjamini-Hochberg FDR = 0.05 (double-sided coefficient *p* value) **c,e** *β*/ORs of *covA* for a specific ancestry in a model including it together with socioeconomic covariates. Independent models are run for different *covA*s; colors label the probed ancestry. **d,f** *β*/ORs of first three *covA* PCs in a model including them together with socioeconomic covariates. The legend also describes an interpretation of the PCs.

As shown above, *covA* exhibits a high correlation across ancestries. Thus we avoided implementing a model with largely multicollinear predictors including *covA* for all ancestries and instead adopted separate models for each ancestry, complementing them with a regression on *covA* PCs (Figure 2b). While *covA*s (Figure 2c,e) highlight the overall excess or lack of certain ancestries in relation with a given phenotype but are largely intertwined, PCs (Figure 2d,f) can be interpreted as independent axes defined by 2 or 3 *covA*s (Figure 2b). Being independent variables in a comprehensive predictive model, they provide a clue to disentangle the potentially collinear *covA* signal and can be reliably used to evaluate significance. When applying this approach, at least one *covA*-based PC had a significant coefficient (coefficient *p* value significant at Benjamini-Hochberg FDR=0.05) in the 16 traits shown in Figure 2c-f out of 27 tested. Furthermore, it is also visible how WHG and Yamnaya tend to be linked with the phenotypic ranges in a similar fashion. As an example, the genomes of taller individuals tend to be more similar to WHG and Yamnaya, while the opposite is true for Anatolia_N and Siberia. PCs are largely consistent with this result, with the PC discriminating Yamnaya and WHG lacking significance.

As we are not controlling for genotype-based PCs in order not to hide potential genome-wide signals, we run the risk of obtaining spurious ancestry/trait associations not caused by genetics. This is due to uneven ancestry similarity across Estonia concurrent with geographically associated socio-economic differences that can influence a trait. Even if such risk is reduced by the extensive sampling of a relatively uniform population, small differences tied to historical reasons^24^ are still visible in *covA* (see Figure 1c,d, Figure S2). Therefore, we include a city/countryside residency covariate in the models, defined as 1 for people living in Tallinn’s county (the wealthiest and most populous) and 0 otherwise, and a covariate for educational attainment, which is a good proxy for family socioeconomic status^25,26^. This control allows us to suggest a significant influence of genomic ancestry on the 16 traits in Figure 2c-f, even when geographical and social stratification is present.

### 2.3 Phenotype-associated genomic regions show specific similarity patterns

To narrow down the signal emerging from the genome-wide analyses we defined three sets of candidate regions by considering windows of 5kb, 50kb or 500kb centered around GWAS catalog^22^ hits for appropriate categories (see Methods and Table S3). As shown in Figure S3, these genomic regions harbor a higher heritability intensity (*h*^2^/Mb) than the whole genome, supporting their appropriateness as candidate regions for the traits of interest.

We then asked whether candidate regions for a given trait showed significantly different coefficients when compared to 50 size-matching random genomic sets, and found it true in 11 out of 27 traits(double-sided Z-test, Benjamini-Hochberg FDR = 0.05), see Z-scores in Figure 3. This analysis has the advantage of naturally controlling for all potential confounders that apply to the genome in its entirety, e.g. social, economic and cultural statuses as introduced in the previous section, thus allowing us to not include any covariates. In addition, this analysis pinpoints genetic signals that are likely to be functionally connected to the trait. Among others, blood cholesterol levels are shown to be positively correlated with similarity to Yamnaya in cholesterol-associated regions with respect to the rest of the genome, while the opposite is true for WHG.

**Figure 3:**
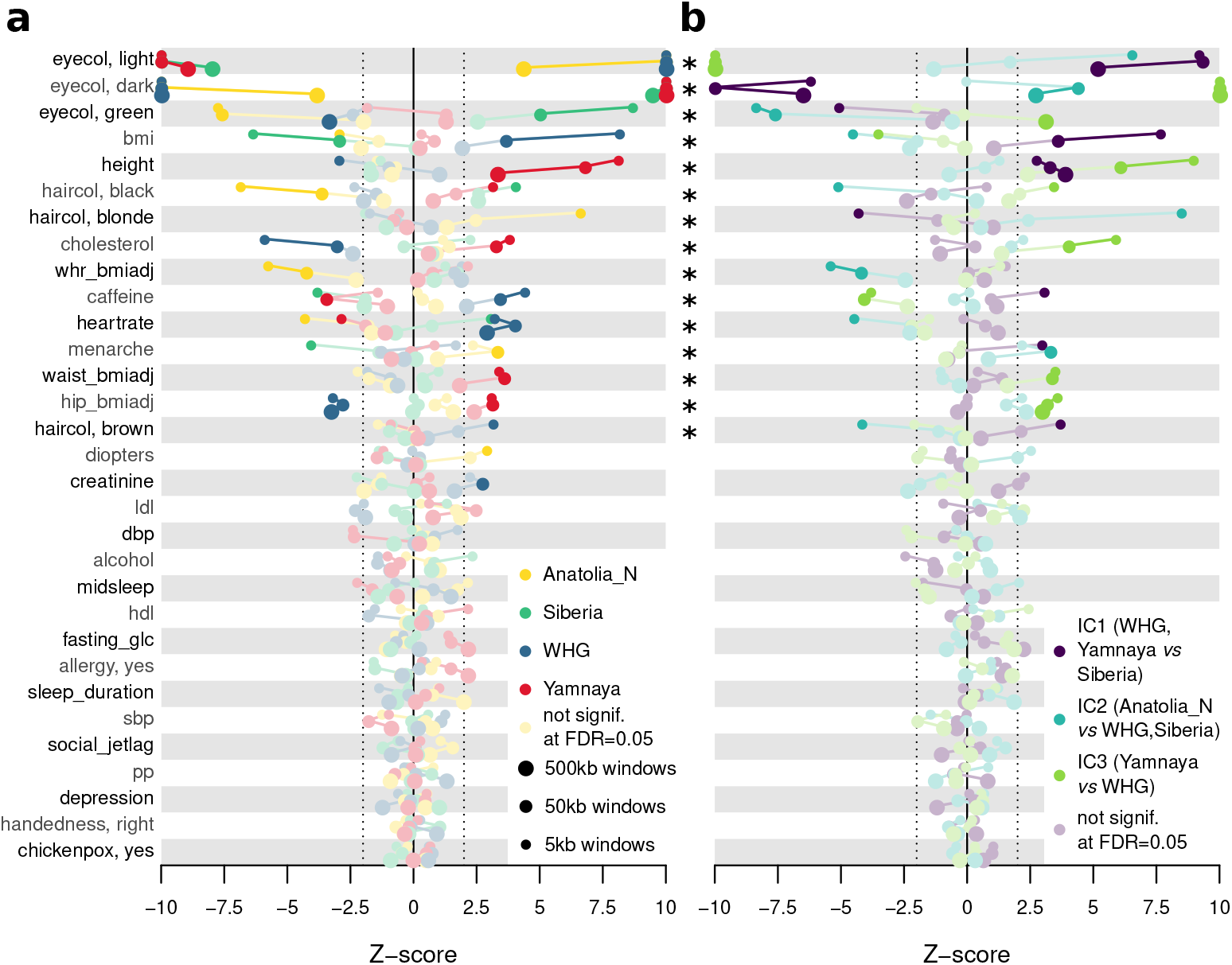
Ancestry-trait association on candidate regions. **a** Z-scores of *covA* coefficients, the color refers to the ancestry probed. **b** Z-scores of coefficients associated with *covA* independent components (IC) computed with whole genome-based *covA* PC loadings. Each color is associated with one of the three ICs. For each trait we show the Z-score of the standardized coefficient associated with candidate regions against a distribution of 50 random genomic regions of matching size. Candidate regions are determined around GWAS hits for appropriate traits as windows with three different widths: 5 (small dot), 50 (medium dot) and 500 (large dot) kilobases. Pastel dots are deemed not significant at Benjamini-Hochberg FDR = 0.05, *p* value from double-sided Z-test; asterisks mark traits to be considered significant according to **b**; dotted lines correspond to absolute Z-scores = 2.

Again, to better interpret the signal and avoid multicollinearity, we transformed *covA*s with the loadings yielded by the PC analysis on whole genome *covA*s (Figure 2b). This, though not returning actual PCs in each candidate region, drastically reduces the collinearity (highest Variance Inflation Factor=1.62 in hair color 50kb candidate regions), while allowing simpler interpretation and, crucially, cross-region comparisons required for Z-scores computation. Indeed this analysis confirms the significance of the association between cholesterol levels and the Yamnaya-WHG axis previously mentioned. In contrast to our genome-wide results, candidate regions no longer yield concordance between WHG and Yamnaya trends across the traits spectrum, both when considering *covA* and their independent components (IC), suggesting a higher specificity of this refined approach.

### 2.4 Selection signatures at candidate regions with ancestry/trait association

So far we only explored associations between a given trait and a local excess of a given ancestry. The observed local admixture unbalance points to a role of that ancient contribution in explaining a given phenotype. However, these results alone do not show whether after the admixture event the incoming genetic material also underwent a selective sweep within the recipient population, altering population-wide allele frequencies as investigated in Mathieson et al. [5]. In other words, the local admixture imbalances we detected so far are not necessarily transferred to the whole population.

We therefore independently asked whether the phenotypes that showed differential contribution from different ancestries exhibit signs of recent natural selection. We applied CLUES^27^ to the list of GWAS hits used above as index for our candidate regions to obtain per-SNP evidence of recent (up to 500 generations ago) natural selection, and to see which phenotypes show enrichment in SNPs with strong selection signals compared to a random set of GWAS hits. Out of the genomic regions responsible for ancestry/trait association shown in Figure 3, pigmentation-related SNPs (eye and hair color) showed extremely high CLUES logLR values (Figures 4a, S4) in accordance with previous results^5,8,28^, as well as SNPs related to BMI and cholesterol, pointing to ongoing or recent selection at these loci. Diastolic blood pressure (DBP) and sleep-related SNPs also showed the same extreme signature, but the candidate regions encompassing them did not reach significance in ancestry/trait association.

**Figure 4:**
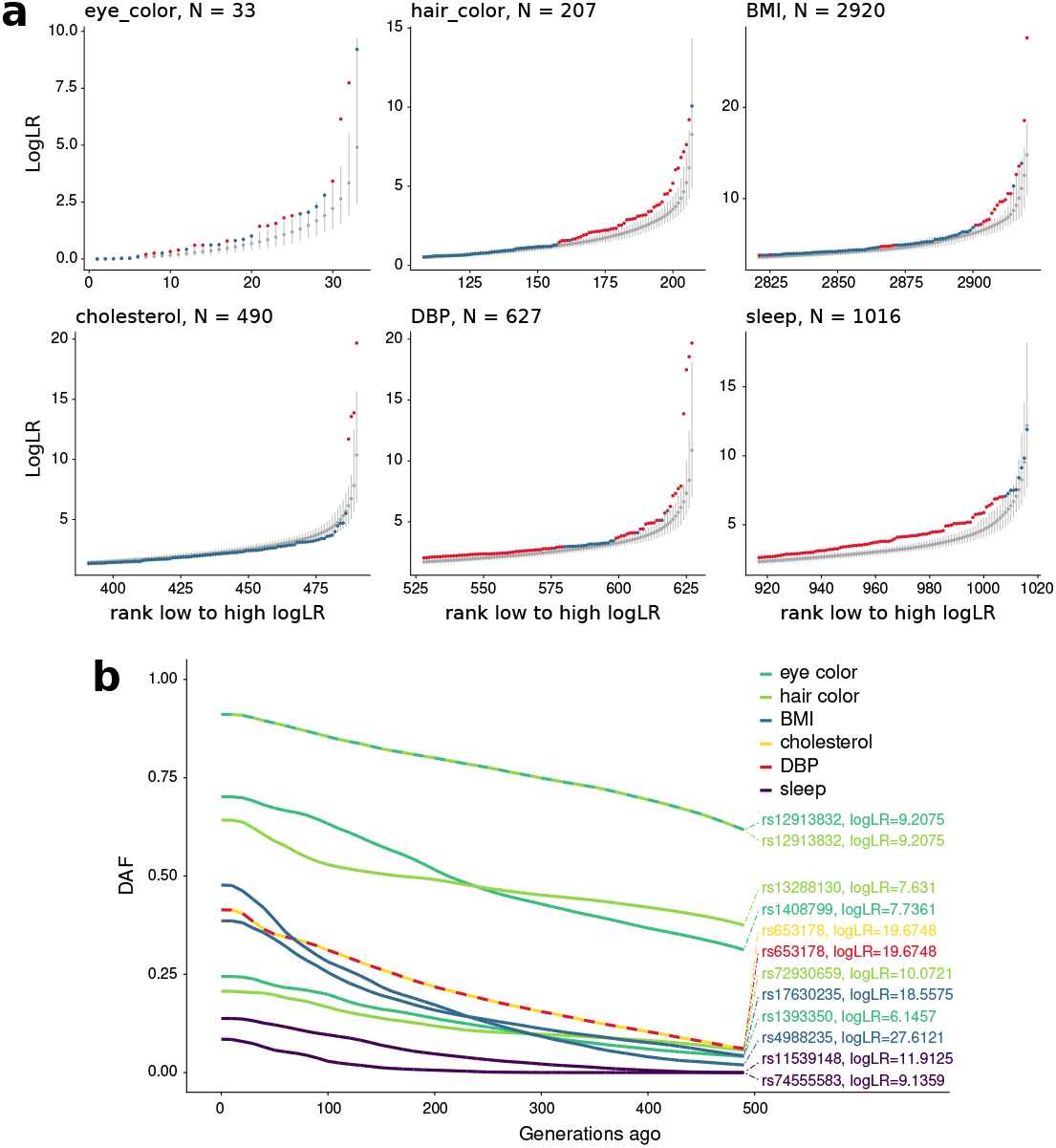
Selection signatures. **a** CLUES log likelihood ratios (logLR) values distribution for GWAS hits for six selected phenotypes. For each phenotype at most 100 top SNPs with highest logLR values and the corresponding ranks from the random GWAS hits distribution are shown. Grey dots show mean values for each rank in the background distribution while the whiskers show the 5-95 percentile range. The logLR values for tested SNPs are shown in red or blue depending on whether the value lies above the 95th percentile of the values from the background distribution with a given rank. Number of tested SNPs for each phenotype are shown in panel titles. **b** Maximum likelihood estimates of derived allele frequency trajectories for top 3 SNPs with highest logLR values for each phenotype. When more than one SNPs come from the same locus, only the top-scoring SNP is shown.

The recent and putatively ongoing nature of the inferred selective pressure on the six traits shown in Figure 4a is further exemplified by the steep increase in derived allele frequencies over time inferred for the top 3 SNPs of each trait and shown in Figure 4b. These include some loci previously shown to be selected in West Eurasians (rs4988235 at MCM6/LCT^29^, pigmentation-related SNPs at HERC2/OCA2, TYRP1, TYR, TPCN2^8,28,30^, rs653178 at ATXN2^31^) and some other yet to be explored: rs17630235, rs11539148, rs74555583.

## 3 Discussion

Putting together the genome-wide, region-specific and selection results, the emerging picture points to a different role of each ancestry in having contributed to the phenotype landscape of contemporary Europeans. As a whole, the most affected traits include pigmentation and anthropometric traits together with blood cholesterol levels, caffeine consumption, heart rate and age at menarche.

In particular WHG ancestry is linked to lower cholesterol levels, higher BMI and putatively contributed light (but not green) eye color to the contemporary Estonian population. Importantly, these associations stand when carefully considering *covA* ICs and, in addition, loci associated with these features also appear to have undergone selection in Estonians. Secondly, although WHG seems to have an association with hip circumference, caffeine consumption and brown hair pigmentation, these evidences are ambiguous.

An enriched Yamnaya ancestry in the pigmentation candidate regions, in contrast with the genome wide analysis, is linked to dark eye and hair colors, consistently with what inferred from aDNA data from the Baltic region^6^. This ancestry is also linked to a strong build, with high stature (in agreement with previous literature^5,7^) and large hip and waist circumferences, both at genome-wide and region-specific levels, but also high cholesterol concentrations when focusing on candidate regions. The associations of Yamnaya and WHG ancestries to respectively higher and lower cholesterol levels, together with the clues of selection at loci connected to cholesterol and BMI, add a critical element to the knowledge of post-neolithic dietary adaptation^6,32,33^ and might have important health-related implications.

Caffeine consumption, although having significant associations, is difficult to connect to a specific ancestry: Yamnaya ancestry seems to be linked with lower consumption, whereas the direction of Siberia and WHG associations depends on the genomic regions included in the analysis.

An enriched Anatolia_N ancestry in the pigmentation candidate regions has implications opposite to Yamnaya, again in contrast with the genome-wide signal. This recurring localized peculiarity of pigmentation loci possibly reflects selection specific to strong GWAS hits as already seen for skin pigmentation^8^. Notably, Anatolia_N enrichment in trait-related genomic regions is connected with a reduced BMI-corrected waist/hip ratio and heart rate. After considering *covA* ICs, this connection between Anatolia_N and heart rate seems to be the one driving the apparent associations of all other ancestries.

Lastly, the Siberia ancestry is connected with dark eye and hair pigmentation, but also green eye color and lower age at menarche. Again, even if this last trait has ambiguous associations with Anatolia_N and WHG ancestries, *covA* ICs provide a clue to disentangle their interactions in favour of a more robust connection with the Siberia ancestry.

Some ancestry/trait associations that were not considered significant at a genome-wide level, are instead discovered when comparing candidate regions to the rest of the genome, possibly due to the higher sensitivity of this approach. On the other hand, the opposite happens for alcohol consumption, depression, sleep duration, social jetlag, diopters, pulse pressure, creatinine levels. This might be due to a misleading or incomplete tagging of the actual functional regions by the GWAS catalog hits, or to an incomplete correction of socioeconomic and other non-genetic factors. In case of sleep-connected traits and DBP, the reported signal of recent or ongoing selection for loci associated to these phenotypes suggests a yet more complex picture.

A general caveat about significance levels observed in this study is that as we refrain from reducing traits by arbitrary choices, even testing multiple alternatives of the same trait, we expose ourselves to inflated false negatives. Complex traits are often interdependent for biological reasons; therefore, when correcting for multiple testing, this risk is intrinsic to this type of analysis. We deemed it best to acknowledge and control it by avoiding overly stringent multiple testing corrections as Bonferroni. In addition, as highly significant traits tend to have higher heritability, it is likely that our analysis might not have enough statistical power for poorly heritable traits.

Taken together, our results show that the ancient components that form the contemporary European landscape were differentiated enough at a functional level to contribute ancestry-specific signatures on the phenotypic variability displayed by contemporary individuals irrespectively to which target population one may examine. In particular, when looking at Estonians, for 11 out of 27 traits surveyed here we could confirm a significant relationship between presence of a given ancestry in genetic regions associated with a given phenotype and how this is expressed by contemporary individuals. While showing that both autochthonous (WHG) and incoming groups contributed genetic material that shapes the phenotype landscape observed today, we also demonstrated that a subset of these loci further underwent positive selection in the last 500 generations. Although not determining whether the selected alleles (and phenotypes) were predominantly contributed by the autochthonous or incoming groups, by connecting genotypic ancestry and complex traits measured in a large dataset, our results reveal both neutral and adaptive consequences of the post-neolithic admixture events on the European phenotype landscape.

## 4 Methods

### 4.1 Sample selection and ancient European grouping

We used 50,353 sequenced or genotyped individuals from the Estonian Biobank^34^ as contemporary Estonian sampleset. After removing second-degree relatives (pi-hat > 0.25) we obtained a subset of 37,952 individuals and used it as a scaffold to perform a PC Analysis (PCA) with Eigensoft-6.1.4. Other individuals were projected on the same PCA space. Outliers identified in this process (with parameters `numoutlieriter: 5 numoutlierevec: 10 outliersigmathreshold: 6`) were discarded. Samples that on the first round of genome-wide *covA*s were more distant than 8 Interquartile Ranges (IQR) from the upper or lower quartile against any of the ancestries were also discarded, resulting in 49811 individuals included in our sample set. For each trait of interest we first removed individuals with missing data for traits and covariates and subsequently discarded second-degree relatives.

To define ancestral European groups we started from the Allen Ancient DNA Resource (AADR) V44.3 merged with present-day individuals typed on the Human Origins array (see Data Availability section). From this set we defined a manually curated core set for each ancestral group, then performed a PCA on a space defined by modern Eurasian and North African individuals west of Iran (included), where the ancient samples were projected. We expanded these core sets to other individuals from AADR dataset using multi-dimensional ellypses with diameters equal to 3 core set SDs. We used 4 dimensions: the annotated dating and the first 3 PCs generated above. With this process we selected 90 WHG, 92 Anatolia_N, 74 Yamnaya S1. In addition, from the ones available from the same dataset, we took 7 samples as representative of the broader Siberian ancestry, assuming any Siberian individual would be equidistant to the other ancestral European groups: S_Even-3.DG, S_Even-1.DG, S_Even-2.DG, Bur1.SG, Bur2.SG, Kor1.SG and Kor2.SG.

### 4.2 Phenotypes treatment and heritability

Continuous traits were treated as specified in Table S2 and regressed against the covariates according to the same table. Individuals with traits or covariates more distant than 4 IQRs from the upper or lower quartile were considered as outliers and discarded. The heritability was computed using LDAK 5.0^35^. First we computed a kinship matrix with the LDAK-Thin Model: we thinned down SNPs on the non-related sample set defined above with parameters `--window-prune .98 --window-kb 100`, then used `--calc-kins-direct` with the resulting weights and `--power .25`. Finally we estimated heritability using REML solver.

### 4.3 *covA* definition

*covA* is the covariance in allele frequency (*p*) within a contemporary individual *i* (i.e. its allele dosage) with the ancestral group of interest *j*, computed respectively against the allele frequency *p_C_* of the contemporary population *C* and the average frequency *p_A_* in all the *A* ancient groups:

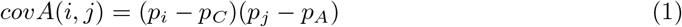

*covA* is expected to be high when the allele frequencies of the individual *i* and the ancestry *j* are similar in comparison with the differences within the contemporary population and across the ancestries that contributed to its genetic makeup. *covA* can be computed across the genome or for specific regions of interest, averaging over the contribution of multiple SNPs. See Supplementary Notes and Figures S5, S6 for further discussion of *covA* properties.

### 4.4 Predicting traits with *covA* and *covA*-based PCs

We fitted each standardized trait with a model including one standardized *covA* and, in case of the genome-wide analysis, socioeconomic covariates as described in the result section. This analysis was restricted to samples for which socioeconomic covariates were defined, i.e. 38,996 samples (including relatives): the actual sample size for this analysis is therefore less than reported in Figure 2a and Table S2. The standardized coefficient (*β* or effect size), or the Odds Ratio (OR) were used to assess ancestry/trait association for continuous and categorical traits respectively. In particular, categorical traits were transformed to {0, 1} where 1 stands for the specified category and 0 for all the others. In addition, each trait was regressed against three *covA*-based PCs, which explained all *covA* variability. PCs were standardized and included together as predictors, socioeconomic variables were again added as covariates. In the candidate regions analysis, we adopted exactly the same steps, performing individual regressions for all the *covA*s and coupling this with a model including all PC-transformed *covA*s. Notably, we transformed all *covA*s using the loadings of the whole genome *covA* PCs, obtaining components that were largely independent, yet not strictly principal. Furthermore, to evaluate association we used coefficient Z-scores computed against the same statistics extracted from 50 random genomic sets with matching size.

### 4.5 Candidate genomic regions

We downloaded GWAS hits from GWAS catalog^22^ (date of download: 20/11/2020) and then extracted for each trait a set of hits connected to it filtering on the reported trait (“TRAIT/DISEASE” field) or selecting the appropriate trait in the Experimental Factor Ontology (EFO) field, as specified in Table S3. Then we took windows of 5, 50 and 500 Kbs centered on the selected hits and merged them where overlapping, obtaining three sets of candidate regions for each trait. To perform the Z-score analysis, for each of them we obtained 50 matching window sets randomly placed across the genome.

### 4.6 Testing for signals of positive selection

In order to test individual SNPs for signatures of positive selection we utilized the Relate/CLUES pipeline^27,36^. This was applied on a curated subset of 1800 unrelated samples; further details on its application are described in Relate/CLUES Supplementary Methods. CLUES was run once for each of the 14,712 unique GWAS hits for traits analyzed here with a derived allele frequency (DAF) above 1% and passing the 1000 Genomes strict mask. To obtain an expected distribution we randomly sampled 10,000 GWAS hits from the GWAS catalog meeting the same conditions and ran CLUES for positions not present among the 14,712 SNPs. Next, for each phenotype we compared its distribution of the logLR values to that of random GWAS hits. We took 1000 random subsets (with replacement) from the 10,000 logLR values each of the same length as the number of GWAS hits for a given phenotype and ranked the logLR values from lowest to highest within each subset. In this way we obtained 1000 values for each logLR rank from 1 to *N* where *N* is the number of SNPs analyzed for a given phenotype. For each rank we calculated the mean and the 5^*th*^ and 95*^th^* percentiles. Finally, we rank SNPs within each trait and compare each logLR value to the mean and 5^*th*^ — 95^*th*^ percentiles range for the corresponding rank of the background distribution. As we are interested in deviations in the higher ranks we focus on the top 100 ranks for each phenotype. Such an approach is conservative as we are testing not against presumably neutral SNPs but against random GWAS hits that are shown to be enriched in signals on natural selection compared to random SNPs in the genome^36^.

## Data availability

The datasets analyzed during the current study are publicly available and can be accessed from the following repositories: data from Estonian Biobank at https://genomics.ut.ee/en/access-biobank (accessed with Approval Number 285/T-13 obtained on 17/09/2018 by the University of Tartu Ethics Committee); AADR plus Human Origins dataset at https://reich.hms.harvard.edu/allen-ancient-dna-resource-aadr-downloadable-genotypes-present-day-and-ancient-dna-data; GWAS catalog at https://www.ebi.ac.uk/gwas/.

## Code Availability

Code for analyses performed in this paper will be accessible upon publication.

## Acknowledgements

This work is supported by the European Union through the European Regional Development Fund, project No. 2014-2020.4.01.16-0024, MOBTT53 (DM, KP, LM, LP); MOBEC008 (VP, MMo, MMe, AE); 2014-2020.4.01.16-0030 (FM, MMe); 2014-2020.4.01.15-0012 (MMe); through the Horizon 2020 research and innovation programme grant no. 810645 (VP, MMo, MMe, AE) and through the Horizon 2020 MSCA Initial Training Network, grant no. 765937 (RC). LS, MMe are supported by the Estonian Research Council through PUT PRG243. SM is supported by the STARS@UNIPD 2019 Consolidator Grant for the project CircadianCare.

## Author Contributions

DM, LP conceived and designed the study; AE contributed in the statistical design; DM, VP performed data analyses; MMo, FM, KP, LV, LM, LP contributed to data analyses; SM, RC provided analyses and expertise about sleep traits; FM, LS, LL, MMe contributed with ancient genetics expertise; DM, LP drafted the manuscript; all authors reviewed and approved the submitted paper.

