## Supplementary Information for "Ancestral contributions to contemporary European complex traits"

Davide Marnetto *et al.*

### 1 Supplementary Notes

#### 1.1 *covA* in simulated ancestry-trait associations

To corroborate the utility of *covA* towards our aim we simulated a modern population composed of three ancestral groups, which separated 10kya and admixed 3kya. We simulated a modern population of 5000 individuals with the following composition: 50% EUR1, 30% EUR2, 20% EUR3. These are simulated groups which separated 10kya and admixed 3kya. Each genome consists of 1000 independent loci of 50kb. We then simulated complex traits as following: we extract randomly one causal SNP in the central quartiles of each window, among those with MAF $> 0.1$, and assign standard-distributed effect sizes. We forced an ancestry-trait interaction by enforcing varying correlations between the effect sizes and the causal allele frequency difference between the contributing ancestry $j$ and the other ancestries. The trait directional shift in the ancestry $j$ is described as average trait value for that ancestral group standardized over all ancestral individuals ($\bar{Z}_j$). The correlation between *covA* for a specific ancestry and phenotype in the modern population correctly indicates that such trait is distinctive of that ancestry as shown in Figure S5a.

#### 1.2 Relations between *covA* and *f*-statistics

We computed *covA* for all individuals in our Estonian sample set against the four ancestral groups as defined in the main text. When comparing *covA* with outgroup $f_3(i, j; Yoruba)$[1], where $j$ is one of the four ancestral groups, the statistics are different but strongly correlated (see Figure S6): this is expected when the $f_3$ outgroup population is an outlier to all populations, contemporary and ancestral, considered in *covA*, as in $f_3(i, j; Yoruba)$. Indeed $covA(i, j)$ has a strong relationship with $f$-statistics[2], i.e. $covA(i, j) = f_4(i, C; j, A) = f_3(i, j, A) - f_3(C, j, A)$ where $C$ is the contemporary population (Estonians in our case) and $A$ is an ideal population with $p = p_A$. Nevertheless, as opposed to $f$-statistics, which only include allele frequencies in actual populations, $covA(i, j)$ has no interpretation in terms of branch length because of the fictitious nature of $p_A$, an allele frequency which only serves as balanced comparison for the ancestries under analysis. In relation with our aim, this constitutes an advantage of *covA*, which does not take into account drift or selection occurred in the branch that connects the outgroup population with the internal node shared by the other populations under analysis. Notably, if we define an outgroup population which separated from all others 125 kya, using $f_3(i, j; outgroup)$ in place of *covA* in the simulated framework proposed above provides less accurate predictions (Figure S5b).

### 2 Supplementary Methods

#### 2.1 Relate/CLUES application

We built local genealogical trees by applying Relate v1.1.4[3] to 2420 phased whole genome sequences of the Estonian Biobank participants described in Kals *et al.* [4] and Pankratov *et al.* [5]. We used the GRCh37_e71 ancestral genome to polarize alleles and applied the strict callability mask for GRCh37 (ref The 1000 Genomes Project Consortium). During the tree-building procedure we used the GRCh37 recombination map[6], mutation rate of 1.25x10-8 and effective population size of 30000. We subsetted 1800 samples to be used for CLUES by removing a) 115 related samples as was done in Pankratov *et al.* [5]; b) PCA outliers (2.5% in both directions along 1st and 2nd PCs both for Estonian-only PCA and when projecting Estonian samples on a "European" PC space); c) singleton count outliers (top/bottom 2.5%); d) samples with pairwise total IBD sharing $\geq$ 166.2 cM with more than one other sample; d) 57 random samples. All analyses used for those filtering steps are described in Pankratov *et al.* [5]. We also randomly selected 100 individuals from these 1800 for coalescence rate through time estimation. We extracted subtrees corresponding to those two subsets using the `SubTreesForSubpopulation` mode of the `RelateExtract` program. We applied Relate's `EstimatePopulationSize.sh` module to the 100 samples described above with mutation rate 1.25x10-8, generation time 28 years, number of iterations 5, tree dropping threshold 0.5 and time bins defined as 10x years ago where x changes from 2 to 7 with an increment of 0.1. We ran CLUES[7] using Relate trees for the 1800 individuals as input as suggested by authors in the GitHub page. We first sampled branch length for the tree containing a SNP of interest (200 samples) by applying the `SampleBranchLengths.sh` module of Relate. We removed 100 of 200 trees as burn-in and selected every 5th tree for importance sampling by CLUES. We focused on the time window between 0 and 500 generations ago.

### 3 Supplementary Tables

**Table S1: Ancestral reference samples.** The table contains for each ancient sample used in the study: ID, ancestral group assignation and if the sample is part of the core set for that group.

**Table S2: Traits analyzed.** The table contains the traits analyzed and how they were encoded, corrected and adjusted for covariates. In addition for each trait we include sample size and case number for categorical traits.

**Table S3: Selection of candidate regions.** The table contains the filter used to select hit SNPs from GWAS catalog for each trait of interest. For each trait we either used a regexp filter on the reported trait field, or selected a specific EFO trait name, as explained in Methods.

### 4 Supplementary Figures

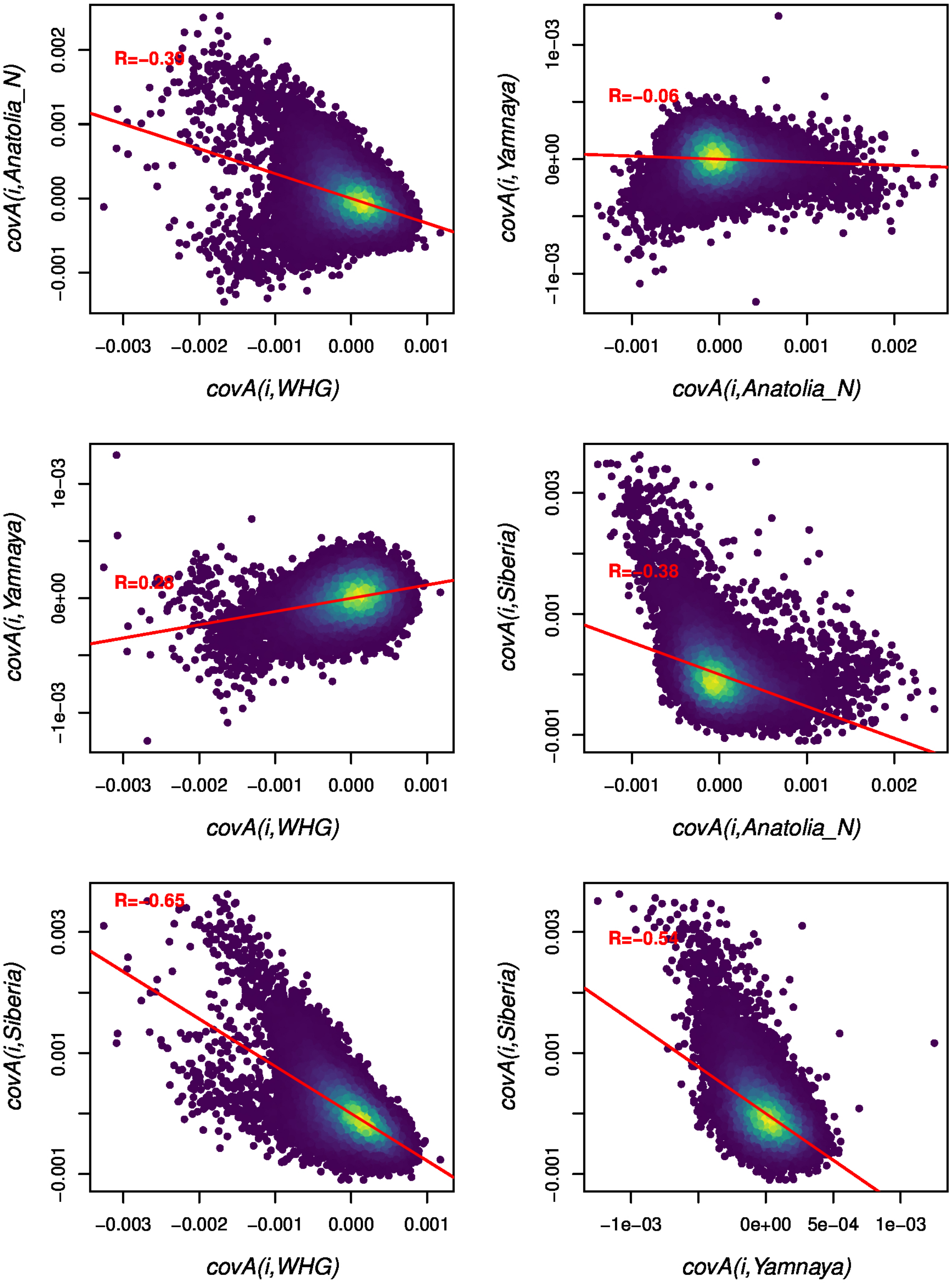


**Figure S1:** ***covA*** **joint distributions for all ancestry combinations.** Each dot is an individual, dots in denser areas are lighter. The red line shows a linear regression, with its R coefficient.

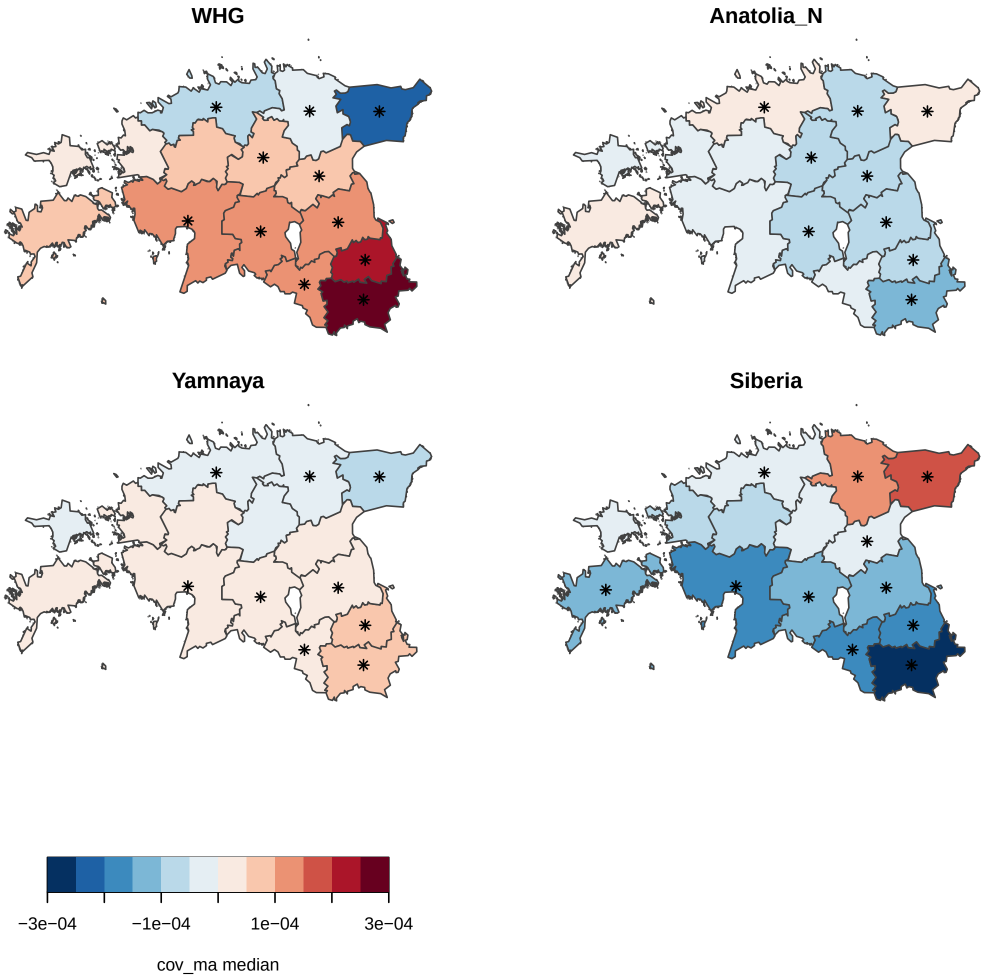


**Figure S2: *covA* distribution across Estonian counties.** Color indicates median *covA* computed in each county while asterisk indicate those counties for which the *covA* is significantly different than the rest of Estonia (two-tailed Wilcoxon-Mann-Whitney test, $p \leq 0.001$)

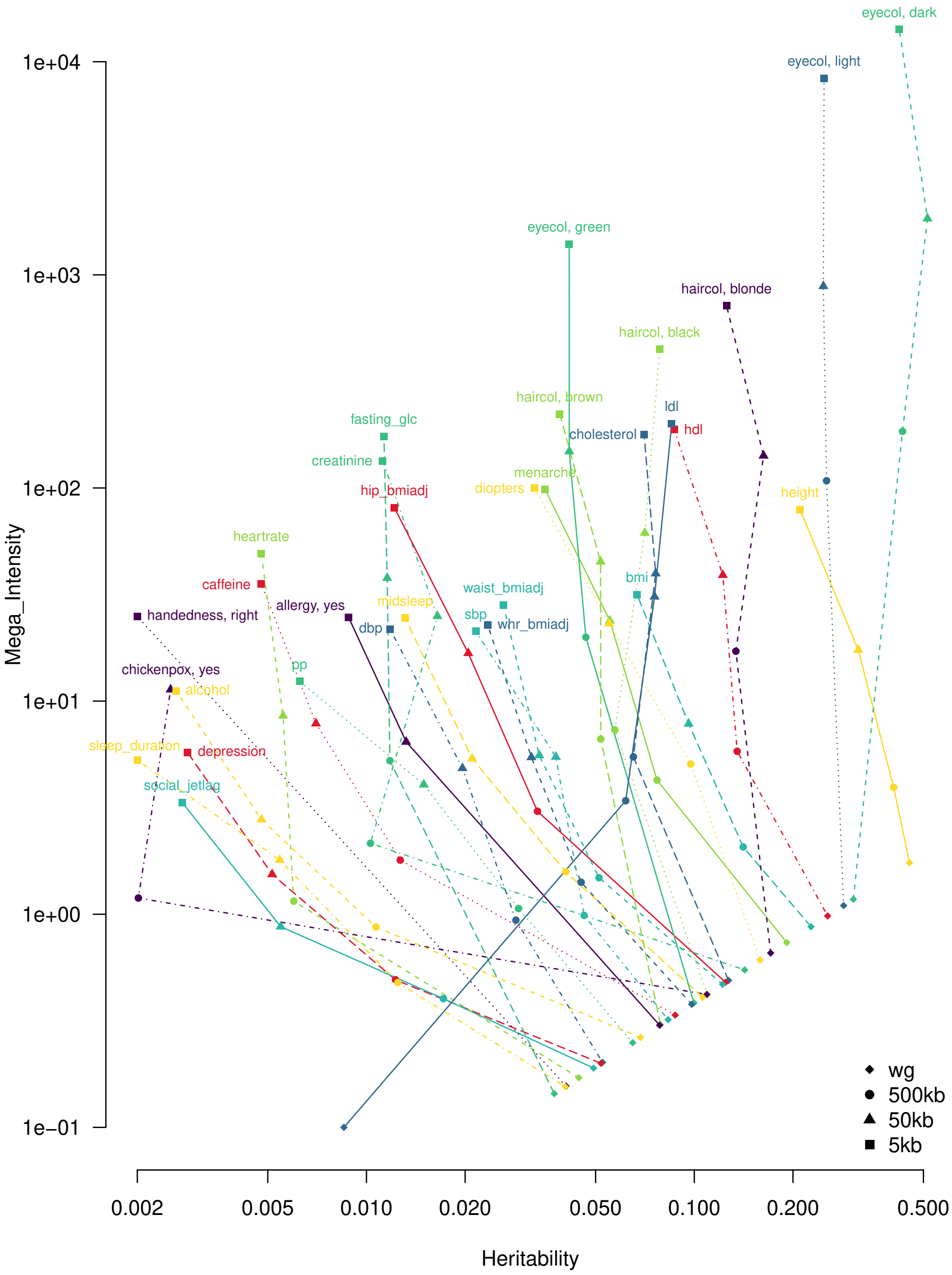


**Figure S3: Heritability of the analyzed traits, for different genomic regions.** Each trait is indicated by a color and a line type, associated with its name,while the point shapes specify which genomic region has been used to compute the heritability, shown on the x axis. The heritability intensity ($h^2$/Mb) is shown on the y axis.

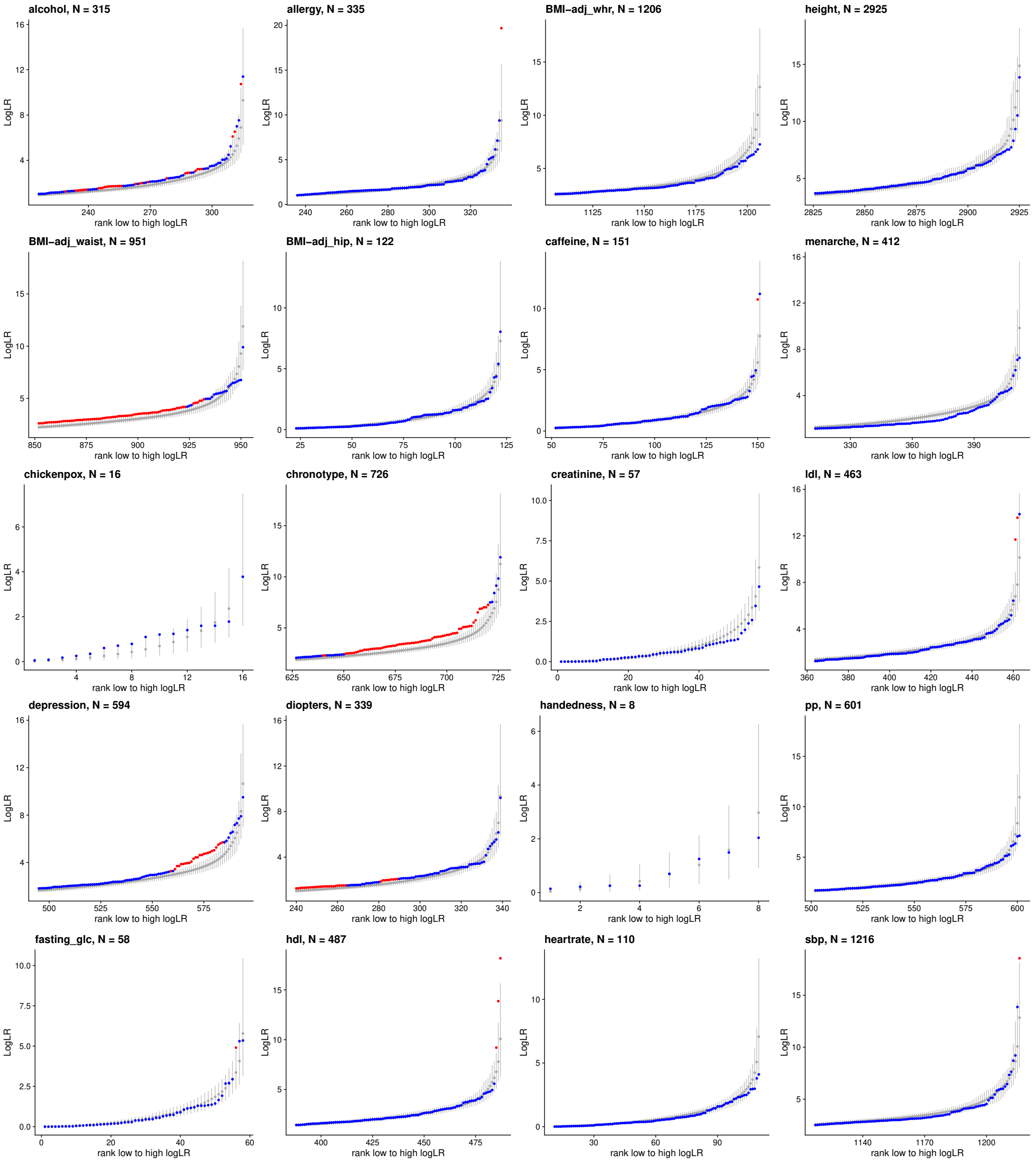


**Figure S4: Selection signatures at GWAS hits for other phenotypes.** CLUES log likelihood ratios (logLR) values distribution for GWAS hits of phenotypes not shown in main Figure 4. For each phenotype at most 100 top SNPs with highest logLR values and the corresponding ranks from the random GWAS hits distribution are shown. Grey dots show mean values for each rank in the background distribution while the whiskers show the 5-95 percentile range. The logLR values for tested SNPs are shown in red or blue depending on whether the value lies above the 95th percentile of the values from the background distribution with a given rank. Number of tested SNPs for each phenotype are shown in panel titles.

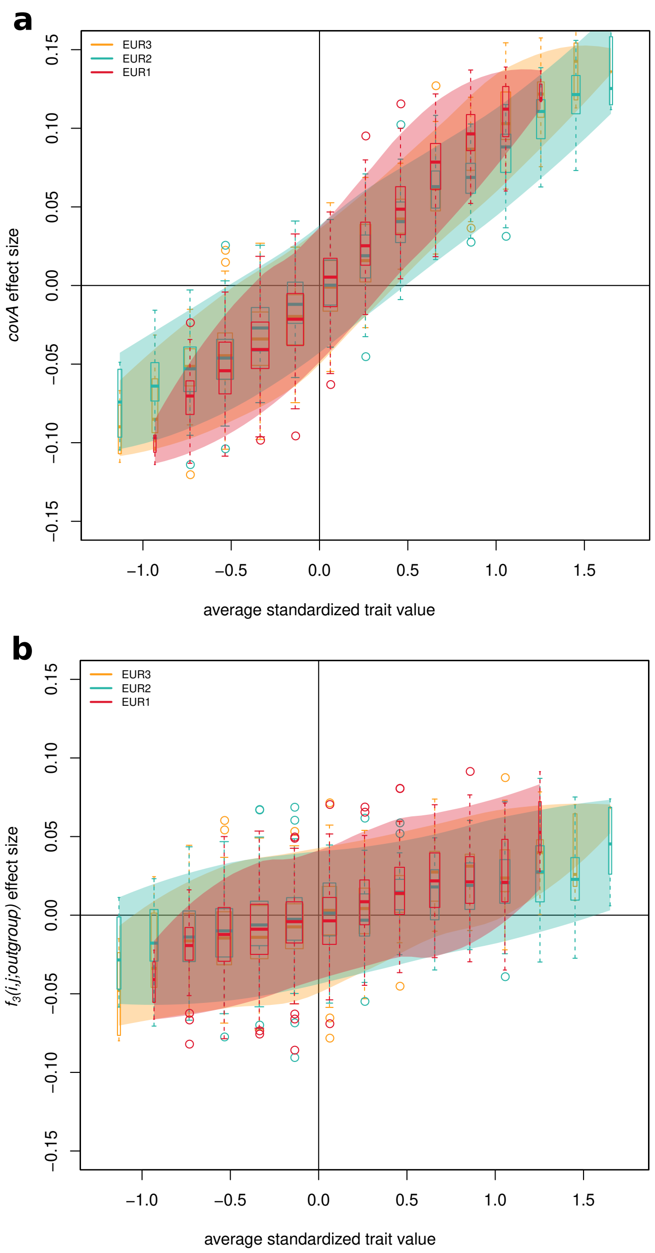


**Figure S5: *covA* and $f_3$ in simulated ancestry-trait associations.** **a** *covA* effect sizes estimated as in Methods section "Predicting traits with *covA* and *covA*-based PCs" for 999 traits variously associated with a contributing ancestry, indicated by color. The shaded area indicates the 95% CI for *covA* effect size at each $\bar{Z}_j$. **b** $f_3(i, j; outgroup)$ effect sizes estimated as for *covA*. The outgroup is a population which separated from all ancestral groups 125 kya.

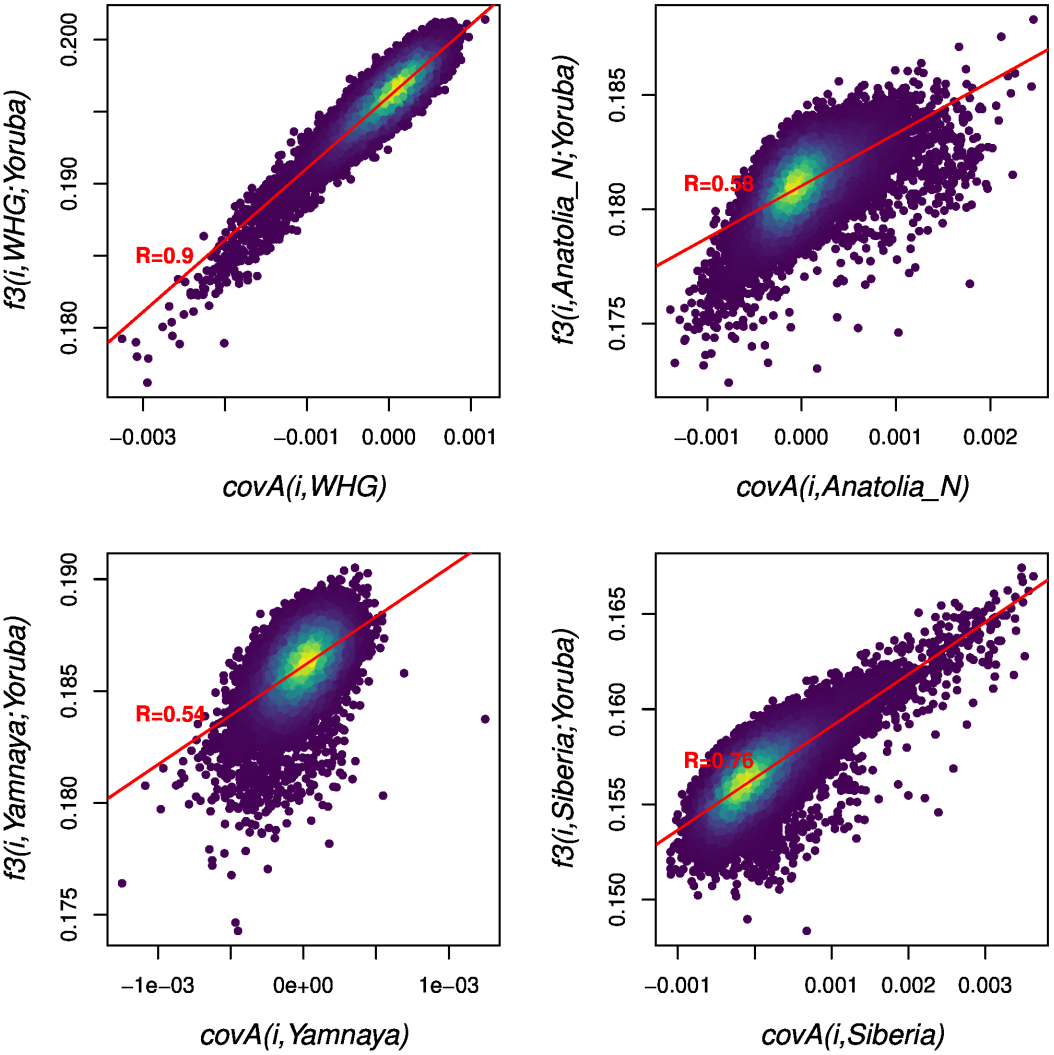


**Figure S6: *covA* and $f_3$ *(i,j;Yoruba)* comparison for each of the four ancestries.** Each dot is an individual, dots in denser areas are lighter. The red line shows a linear regression, with its R coefficient.
